# HIV-associated non-Hodgkin lymphoma tumor-microenvironment axes differ by EBV status across cellular origins

**DOI:** 10.1101/2025.10.15.682084

**Authors:** Amy Chadburn, Maria Montserrat Aguilar Hernandez, Joanne Dai, Mitra Harrison, Nicolas M. Reinoso-Vizcaino, Ashley P. Barry, Emily Hocke, Karen Abramson, Vaibhav Jain, Cliburn Chan, Ethel Cesarman, Micah A. Luftig, Elliott D. SoRelle

## Abstract

Non-Hodgkin Lymphoma (NHL) is the main cancer-related mortality for people living with HIV (PLWH). NHL genetic and molecular classifications have been intensely studied and correlated with clinical outcomes, but critical unanswered questions relevant to malignancy persist. For example, tumors positive for Epstein-Barr virus (EBV) are aggressive and account for 30-50% of HIV-associated NHLs, yet insights on the nature of EBV in NHL pathogenesis or potential therapeutic vulnerabilities have been limited. Here, we examined HIV-associated NHLs stratified by histopathologic classification, cell of origin (COO), molecular subtype, and EBV status using genome-wide spatial transcriptomic analyses. Tumor tissues in EBV^+^ HIV-NHLs displayed enriched expression of mitochondrial respiration and purine metabolism signatures versus EBV^-^ tumors. Immune infiltration of EBV^+^ tumors was limited, and immune and stromal regions of EBV^+^ HIV-NHLs displayed upregulation of the gene for the iron exporter ferroportin (*SLC40A1*), indicating immunosuppressive macrophage polarization. An immunosuppressive *SPP1-CD44* axis and oncogenic *GAS6-AXL* interaction between tumor and stroma were specifically predicted for EBV^+^ HIV-NHLs with plasmablastic features. Tumor microenvironment (TME) similarities to secondary lymphoid tissues such as inverse *CXCL12-CXCL13* gradients and evidence of matching between tumor B cell phenotypes and reticular cell signatures were also observed. Finally, we developed a simple yet flexible approach to quantify expression gradients and cell proximity to annotated regions. Thus, this study highlights potential avenues for NHL therapy tailored by EBV status and provides a unique resource to examine tumor-TME interactions in the HIV-immunocompromised context.

**Key Points:**

- Spatial transcriptomic landscapes resolve cell-of-origin- and EBV-associated HIV-NHL heterogeneity including immunosuppressive myeloid responses to EBV+ tumors
- Spatial gradients including CXCL12 and CXCL13 indicate that tumor-stromal interactions mimic lymphoid tissue organization and depend on B cell development stage
- Development of a direction-agnostic method to calculate spatial expression gradients relative to annotated tissue and cell types of interest
- Predicted pharmacologic vulnerabilities in clinical samples are replicated *in vitro* and confirmed by small molecule screening

## Introduction

People living with HIV (PLWH) experience elevated risk (∼11x) for lymphoma versus the general population due to compromised T cell-mediated control of lymphoproliferation^1^. Lymphoma remains the primary cancer-related mortality for PLWH in the antiretroviral therapy (ART) era^2^. Most HIV-related lymphomas (80%) are non-Hodgkin lymphomas (NHL) including diffuse large B cell (DLBCL), plasmablastic (PBL)^3^, and Burkitt Lymphomas (BL)^4^ as well as polymorphic B-lymphoproliferative disorders (B-LPD)^5^. NHLs are delineated by histopathologic cell-of-origin (COO), immunohistochemistry (IHC)^6^, genetic profiles^7-10^, location, and whether tumors are Epstein-Barr virus (EBV) positive^11,12^. COO and genetic classifications impact prognosis and treatment and have been studied intensively through microarrays and sequencing^7,8^. Spatial interactions between NHL microenvironments and tumors are likewise important for classifying disease heterogeneity^13,14^ but remain incompletely understood. For example, B cells in different developmental stages interact with distinct immune and reticular cell milieus in lymphoid tissue^15,16^. Whether such differences are recapitulated – or dysregulated – in tumor-TME interactions dependent on COO is unclear. High-dimensional *in situ* assays are suited to address such questions and implications for pathogenesis.

PLWH are more likely to develop EBV-associated lymphomas, even with ART. Indeed, 30-50% of HIV-NHLs are EBV-positive^17^, far exceeding the general population frequency (∼5%)^18,19^. In the Germinal Center (GC) model, EBV infects naïve B cells and enacts coordinated latency programs (Latency IIb, III, IIa, I) to mimic B cell activation and differentiation^20,21^ and establish persistence in memory B cells^22,23^. The GC model was founded on two clinical observations: lymphomas originating from most mature B cell stages can be EBV^+^, and patterns of EBV nuclear antigens (EBNAs) and latent membrane proteins (LMPs) vary by COO^24^. GC-derived EBV^+^ BL is classically Latency I (EBNA1^+^); post-GC Hodgkin Lymphoma is Latency IIa (EBNA1^+^EBNA2^+^LMP1^+^); and DLBCLs may display several profiles depending on GCB versus ABC origins^25,26^. However, infection profiles only provide shorthand diagnostics since viral expression is heterogeneous^27^. For example, primary B cells infected with EBV *in vitro* exhibit a phenotypic spectrum within 1-2 weeks that includes activated precursor B cells (AP, pre-latent), proliferating centroblast-like B cells (B prolif, latency IIb), centrocyte-like B cells with high NF-κB activity (NF-κB Act, latency III), and post-GC plasmablast-like cells (PB)^28^. This continuum is retained in EBV-immortalized lymphoblastoid cell lines (LCLs)^29^, exhibits oscillatory plasticity reminiscent of a model for ABC-DLBCL phenotypic complexity^30^, and includes cells with autoimmune-associated features found in DLBCL precursor cells^31,32^. Such variation may affect tumors because many EBV genes engage B cell programming, modulate antiviral responses^33^, and induce potent immunity^34^. Notably, EBV status currently does not impact NHL therapies^18,35^. Given the aggressive course and overrepresentation of EBV^+^ NHLs in PLWH, deeper understanding of EBV-specific HIV-NHL pathobiology is needed to improve therapies and reduce health disparities.

We performed high-dimensional *in situ* HIV-NHL characterization with spatial transcriptomics (ST), ISH, and IHC to determine whether HIV-NHL composition is COO- and/or EBV-dependent. Based on observed concordance between EBV^+^ PBL and single-cell RNA-seq (scRNA-seq) phenotypes of EBV-transformed LCLs, we further explored potential EBV^+^ PBL pharmacologic vulnerabilities via inhibitor screening in LCLs. Cell-cell communication and distance statistics were also applied to analyze COO- and EBV-stratified tumor-TME interactions. This approach yielded unique resources to support HIV-NHL research and exploit EBV-dependent vulnerabilities.

## Methods

### Samples

Ten FFPE HIV-NHL samples (**Table S1**) were collected (residual after diagnosis) from New York Presbyterian Hospital–Weill Cornell Medicine Department of Pathology and Laboratory Medicine with IRB approval. Research was conducted in accordance with the Declaration of Helsinki. Diagnoses were made using World Health Organization criteria^36^, histomorphology, and immunohistochemistry. GCB versus non-GCB DLBCL subtypes were determined via Hans algorithm^6^. 5μm serial sections were produced for ST and ISH. 10μ;m sections were used for quality control (QC).

### IHC/ISH

Monoclonal antibodies were used against: CD10 (56C6; Leica Microsystems, Wetzlar, Germany), BCL-2 (124), BCL-6 (PG-B6p), MUM-1/IRF4 (MUM1p) and Ki-67 (MIB-1) (DakoCytomation, Carpinteria, CA). EBV Probe ISH Kit (Leica Microsystems; Vision BioSystems Novocastra, Newcastle-upon-Tyne, UK) was used for EBV RNA (EBER) *in situ* hybridization (ISH). Cases were considered positive for CD10, BCL6, and IRF4 when >30% of neoplastic cells were immunoreactive; BCL2 positivity was defined when ≥50% of cells had moderate/strong positivity. Nuclear Ki-67 was determined semi-quantitatively as positive % of tumor cells. Cases were considered EBER-positive when hybridization signal was identified in most neoplastic cells. EBV latency and lytic replication were evaluated by IHC with antibodies to LMP1 (CS1-4, Abcam, Cambridge, UK), EBNA2 (PE2; Dako), LMP2A (15F9; Abcam), and Zta/ZEBRA.

### ST preparation & sequencing

RNA quality from tissues on charged slides was assessed by DV200 (% of fragments >200nt). Curls were scraped, deparaffinized, and RNA-extracted via Qiagen RNeasy Kit (Cat# 73504). Fragment distributions were analyzed on an Agilent TapeStation 4200 using High Sensitivity RNA ScreenTape and reagents (**Table S2**). Serial 5μ;m FFPE sections were mounted, incubated at 42°C for 3h, and stored at RT in a desiccator for <2w. Samples were deparaffinized and H&E stained according to 10x Genomics Visium CytAssist protocol CG000520. Slides were brightfield imaged (Zeiss AxioScan Z1) at 20x. Slides were de-stained, de-crosslinked and processed using 10x Genomics Visium CytAssist Spatial Gene Expression Reagent Kit (Cat# 1000520/1000522; protocol CG000495). Tissues were assembled into CytAssist cassettes and reacted with human whole transcriptome probes for hybridization and ligation. Tissue slides and Visium CytAssist Spatial Gene Expression Slides were loaded in the CytAssist instrument for library transfer. Probes were released from tissue and captured by oligonucleotide-barcoded spots on Visium slides (55μ;m spot diameter). Visium slides were removed and used to prepare libraries, which were analyzed via Agilent TapeStation. Libraries with median length ≈260nt passed QC and were sequenced at 40,000 target depth (Illumina Novaseq 6000 S2 2x50 paired end flowcell with 28bp read 1, 10bp i5 and i7 indices, and 50bp read 2).

### Bioinformatics

Bases were assembled into reads, aligned against the human genome (hg38) to generate UMI count matrices, QC filtered, and analyzed as an annotated, integrated object containing all HIV-NHLs using standard software tools and single-cell references^29,37-42^ (see **Supplementary Text, Files S1-S2**)

### Inhibitor screening

LCL viability was screened after treatment with a Selleck Bioactives 2,100 compound library (Selleck Chemicals LLC, Houston, TX) with functionally annotated NMR and HPLC-validated FDA-approved drugs, natural products, and chemotherapeutics. Cells in 384-well (400/well) or 96-well (1500/well) plates were treated (10μ;M compounds) for 72 h, and viability versus DMSO-treated LCLs was evaluated via CellTiter Glo (Promega). Initial screening was performed with single replicates; candidates were screened subsequently in triplicate. Hits were determined by significant concordance in wild-type LCLs and dCas9-KRAB-transfected LCLs lacking sgRNAs. 10 μ;M SMI-16a (MedChemExpress, cat. #HY101947) testing was performed in Ly3, Pfeiffer, Farage, IBL1, and LCLs and quantified as described above.

### Data

ST data are deposited in NIH GEO (GSE274051).

## Results

### HIV-NHL samples and spatial transcriptomic (ST) characterization

Tissue from ten HIV-NHLs (5 EBV^+^, 5 EBV^-^) of various COO (**Table 1**) were analyzed using Visium V2 spatial transcriptomics (ST) with CytAssist (**Figure 1A-B**). ST sections were cut adjacent to EBER-stained sections to ensure spatial correspondence. Samples were chosen across a phenotypic spectrum to capture HIV-NHL diversity and investigate features independently associated with EBV. Molecular classifications of EBV-associated lymphomas in PLWH remain to be determined since EBV^+^ cases typically have fewer mutations within NHL subtype and conform poorly to existing subgroups^43,44^. An HIV-associated EBV^+^ polymorphic B-LPD was included, as this disease is undercharacterized^45,46^. Integrated ST datasets exhibited highly correlated genome-wide expression (Pearson R=0.905) and global alignment between EBV^+^ and EBV^-^ samples (**Figure 1C**). Overall, 60,481 spots were analyzed (42.9% from EBV^+^, 57.1% from EBV^-^ samples). ST data were stratified and analyzed by sample, COO, and EBV using Seurat^37,38^ (**Figure S1**). Proliferation scores were calculated from S- and G2M-phase genes; for EBV^+^ samples, proliferation correlated to EBER^+^ regions (**Figure 1D**). Coarse tissue annotations (tumor, immune, stroma) were determined by grouping high-resolution phenotypes found via unsupervised methods (**Figures 1E, S2**). ST phenotypes in tumor tissue were consistent with protein IHC molecular classification^6^ (**Figure 1F**). *BCL6* and *CD10 (MME)* expression was observed in GC-derived tumors (n=3), although *BCL6* was reduced in EBV^+^ BL, consistent with virus-mediated downregulation^47,48^. Conversely, *IRF4* was elevated in NHLs of non-GC COO, particularly EBV^+^ tumors. Overall, HIV-NHL tumors from GC and indeterminate (IND) COO expressed B lineage markers (*CD19, CD20/MS4A1, CD22, CD79A*) versus non-GC COO tumors with plasmablast signatures (*XBP1, MZB1, BLIMP1/PRDM1, CD38, CD138/SDC1*).

**Figure 1.**
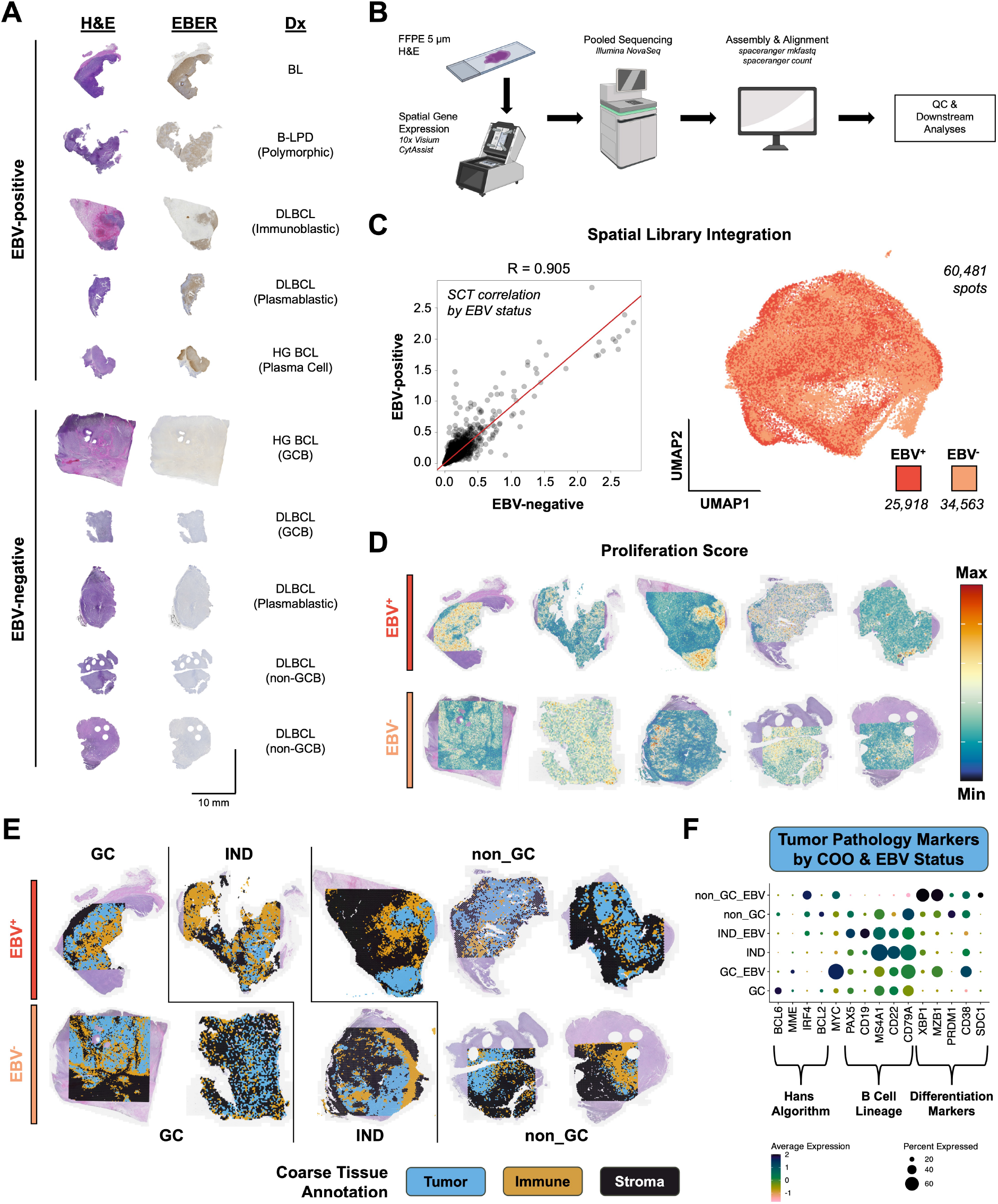
Spatial transcriptomic study of HIV-NHLs. **(A)** H&E and EBER staining of EBV^+^ and EBV^-^ HIV-NHLs analyzed with 10x Visium ST (Dx = diagnosis; BL = Burkitt Lymphoma; B-LPD = B Lymphoproliferative Disease; DLBCL = Diffuse Large B Cell Lymphoma; HG BCL = High Grade B Cell Lymphoma; GCB = Germinal Center B COO). **(B)** Overview of Visium ST workflow with FFPE H&E sections using CytAssist. **(C)** Genome-wide average expression correlation (left panel) and integration (right panel) of all ST datasets, stratified by EBV status. **(D)** Calculated *in situ* proliferation scores in HIV-NHLs by EBV^+^ (top row) and EBV^-^ (bottom row) status. Scores are based on S-phase and G_2_M-phase gene modules and overlaid on H&E images. **(E)** Segmentation of tumor (blue), immune (gold), and stroma (black) regions in EBV^+^ (top row) and EBV^-^ (bottom row) HIV-NHLs. Lines divide samples by tumor COO (GC = germinal center COO; IND = indeterminate COO; non_GC = non-germinal center COO). **(F)** ST expression dotplot of diagnostic biomarkers and B cell lineage genes in tumor regions stratified by COO and EBV status. Dot size denotes the fraction of annotated tumor spots expressing a given gene; dot color denotes relative expression level (SC Transform assay).

### HIV-NHL annotation and expression by COO and EBV

HIV-NHLs were analyzed using differential expression (DE) across tissue types (**Figure 2A**). Multiple intratumoral B cell phenotypes were delineated: a pre-GC activated precursor (AP) state expressing *FCRL5* and *CD22*, actively proliferating cells (B Prolif) expressing *MYBL2, POU2AF1*, and *MCM7*, and differentiated plasmablasts (*MZB1, IRF4, JCHAIN*). Tumor-adjacent immune regions contained CD4^+^ and CD8^+^ T cells but were dominated by macrophages (*CD68, C1QA, C1QB*). Stroma scored highly for fibroblasts, pericytes, and endothelial cells, with subsets of CCL19^+^ fibroblasts (CCL19+ Fib) and ISG15^+^ pericytes and cancer-associated fibroblasts (ISG15+ PCs, ISG15+ CAFs, respectively). HIV-NHLs exhibited variable tumor, immune, and stromal compositions when stratified by COO (**Figure 2B**). Despite limited sample sizes, significant B Prolif enrichment (reflecting GC DZ-like composition) was observed in GC-derived HIV-NHLs (Kolmogorov-Smirnov test, p=0.017), and plasmablasts were enriched in non-GC tumors (p=0.032).

**Figure 2.**
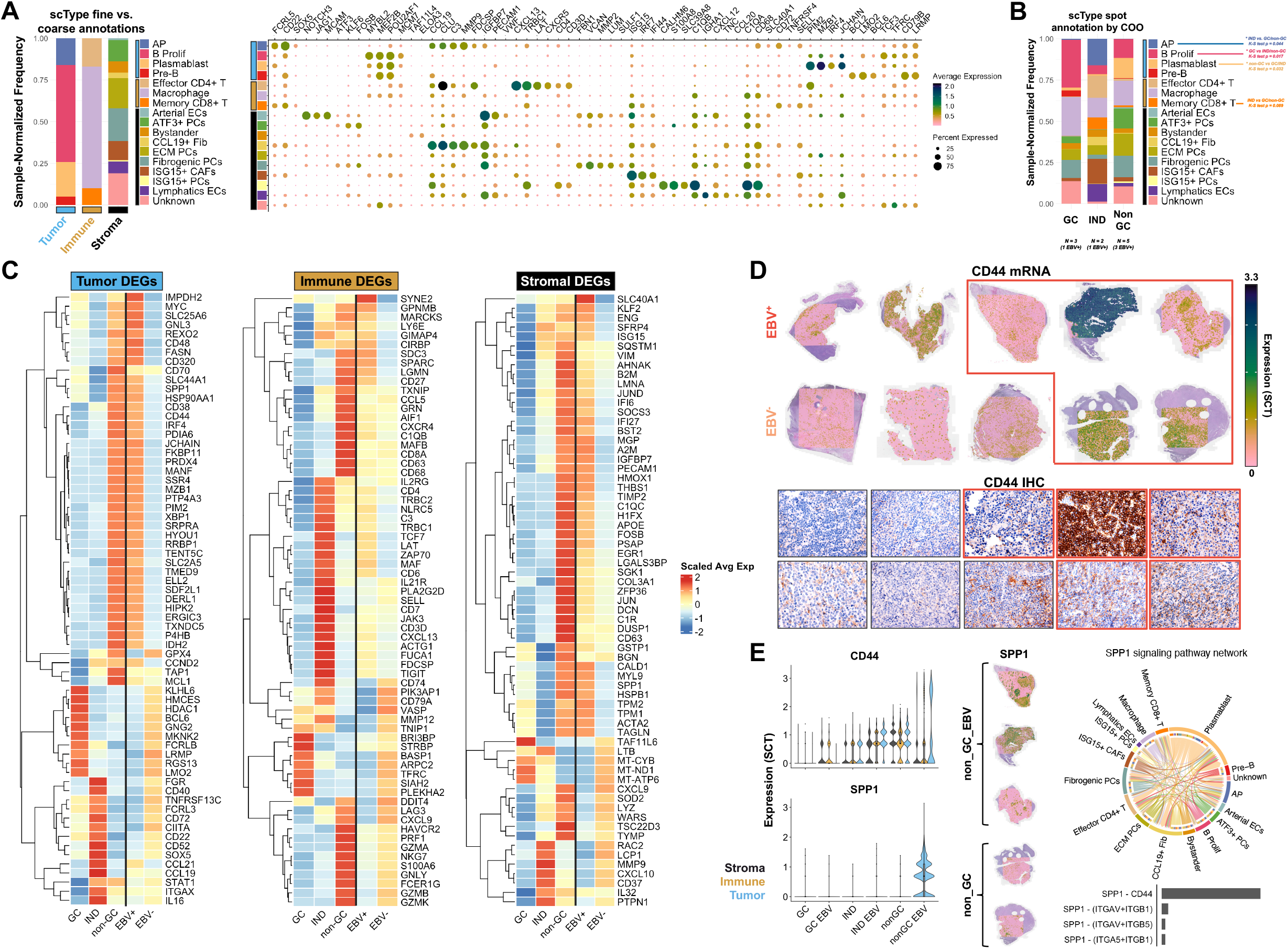
HIV-NHL phenotypic annotation and DE by cell of origin (COO) and EBV status. **(A)** Cell type composition (left panel) and dotplot of expression signatures (right panel) in integrated HIV-NHL ST data. Each coarse tissue type (tumor, immune, stroma) is represented as constituent cell types calculated with scType. **(B)** COO-stratified cell type composition. Cell types are color-coded and grouped by coarse annotation (tumor, immune, stroma) as in (A). Data are from 3 GC-derived tumors (1 EBV^+^), 2 IND-derived tumors (1 EBV^+^), and 5 non-GC tumors (3 EBV^+^). Lines to cell type names denote results of Kolmogorov-Smirnov significance tests comparing cell type frequencies by tumor COO (* indicates p<0.05). **(C)** DE genes by COO and EBV status identified through within-tissue (tumor, immune, stroma) comparisons. Scaled average expression is presented. Within each tissue type, DE by COO is presented in three columns (GC, IND, non-GC) to the left of the vertical black line and DE by EBV status (EBV^+^, EBV^-^) is presented in two columns to the right of the vertical black line. **(D)** ST expression of CD44 mRNA (top panel) and IHC protein (bottom panel) in HIV-NHLs. EBV^+^ samples are in the top row of each panel, and all non-GC samples outlined in red. **(E)** CD44 and SPP1 (osteopontin) expression by tissue, COO, and EBV status (left panels). Circos plot of predicted SPP1 signaling network enriched in EBV+ non-GC HIV-NHLs (right panel).

DE genes (DEGs) by COO and EBV were identified within coarsely-segmented tissues (**Figure 2C**). Stroma from EBV^+^ samples exhibited elevated interferon-induced signatures (*ISG15, IFI6, IFI27, BST2, C1QC*) and upregulated *SLC40A1* (the iron exporter ferroportin, a marker of immunosuppressive cancer-associated macrophages^49^) across COO subtypes (**Figure 2C**, left). Immune infiltrates in EBV^+^ cases also exhibited distinct characteristics: upregulated *SYNE2* (a marker of *SYNE1*^+^*SYNE2*^+^ T cells in people with EBV^+^ hemophagocytic lymphohistiocytosis and infectious mononucleosis^50^) and evidence of restrained macrophage inflammation (*GPNMB, LY6E*)^51,52^. However, pro-inflammatory *MARCKS* was also enriched in EBV^+^ sample immune regions. In EBV^-^ samples, pro-tumoral expression enrichment included ubiquitin ligase *SIAH2*^53,54^ (only in GC-derived HIV-NHLs) and *MMP12* suggestive of myeloid suppressor function^55^ (GC-derived and IND HIV-NHLs) (**Figure 2C**, middle). Thus, DE analyses suggested COO- and EBV-dependent immunosuppressive microenvironments.

Tumor DEGs were consistent with molecular phenotype and COO (**Figure 2C**, right). Non-GC tumors (especially EBV^+^) had differentiation signatures (*CD38, CD44, IRF4, XBP1, MZB1*) whereas GC-derived tumors retained GC hallmarks (*BCL6, KLHL6, LRMP, RGS13, LMO2*). Signatures for oxidative stress (*PRDX4, MANF, HYOU1, TXNDC5*) and ER stress (*HSP90AA1, PDIA6, FKBP11, ELL2, P4HB, ERGIC3*) were enriched in EBV^+^ non-GC tumors. EBV-specific signatures for nucleotide metabolism (*IMPDH2, GNL3, REXO2*)^56,57^, fatty acid synthesis (*FASN*), mitochondrial ADP/ATP and choline transporters (*SLC25A6, SLC44A1*)^58,59^, and *MYC* were also observed across tumor COO. These signatures imply EBV-dependent metabolic pathways in HIV-NHLs. Transcription factor activity predictions indicated COO-specific regulation and corroborated enhanced MYC activity in EBV^+^ HIV-NHLs (**Figure S3A-B**). Estrogen signaling was associated with EBV in non-GC tumors and MAPK activity was upregulated in EBV^+^ tumors across COO (**Figure S3C**). Hypoxia and

TRAIL networks were downregulated in EBV^+^ versus EBV^-^ tumors (COO-matched comparisons; **Figure S3D**), implying differential resistance to oxidative stress and apoptosis.

Following up on tumor DEGs, we focused on *CD44* enrichment in non-GC tumors (particularly EBV^+^ cases), which was consistent with prior reports and bears prognostic significance^60-63^. CD44 elevation in non-GC tumors was confirmed by IHC (**Figure 2D**). Although CD44 enrichment in non-GC tumors was independent of EBV, intratumoral expression of the CD44 ligand osteopontin (*SPP1*) was specific to EBV^+^ non-GC cases (**Figure 2E**). Cell-cell communication analysis^64^ revealed a *CD44-SPP1* axis enriched in non-GC EBV^+^ HIV-NHLs, and plasmablastic tumors co-expressed *CD44* and *SPP1* suggestive of pro-survival autocrine interactions^65,66^. *CD44* was also observed in CD4^+^ and CD8^+^ T cells, consistent with tumor-immune crosstalk. Microenvironment *SPP1* expression in macrophages and fibroblasts further supported tumor-immune-stromal *CD44-SPP1* activity (**Figure 2E**). These EBV-associated interactions are noteworthy given immunosuppression and oncogenicity of *CD44-SPP1* signaling in other malignancies^67-70^. Multiple CD44-convergent ligand-receptor axes including collagen, fibronectin, and MIF were also conserved independently of EBV status (**Figure S4**).

LCL growth inhibitors target features enriched in EBV^+^ PBL including PIM kinase activity LCL scRNA-seq signatures^29^ were scored across HIV-NHLs to evaluate model correspondence to clinical presentations. EBV^+^ PBL (immunoblastic, plasmablastic, and plasma cell NHL) tumors scored strongly for LCL plasmablast and B Prolif phenotypes (**Figures 2A, 3A, S5**). PBL gene ontologies included oxidative phosphorylation, mitochondrial metabolism, protein folding, and ER stress (**Figure 3B**). Given this correspondence, we investigated LCL vulnerabilities to small molecules. 159 of 2,078 compounds (7.65%) yielded significant LCL killing (**Figure 3C, File S3**), ∼25% of which (39) inhibit biological activities consistent with identified GO terms. Inhibitor targets included the proteasome, protein trafficking, HSP90, mTOR, and purine metabolism. Significant killing by a pan-PIM kinase inhibitor (CX-6258 HCl) was noteworthy given upregulation of *PIM2*, an anti-apoptotic serine-threonine kinase that stabilizes c-MYC^71-73^, in EBV^+^ non-GC HIV-NHLs (**Figure 3D**). Based on DepMap CRISPR perturbation data^74^ indicating *PIM2* selectivity in mature B cell neoplasms (**Figure 3E**) and efficacy of PIM2 inhibition for multiple myeloma^75-77^, we treated EBV^-^ (Ly3, Pfeiffer) and EBV^+^ (Farage, IBL1, LCL) lines with PIM2-selective SMI-16a. EBV^+^ lines were significantly more sensitive to SMI-16a (**Figure 3F**; p=0.039, two-tailed t-test). Thus, LCLs may recapitulate certain HIV-NHL characteristics and EBV^+^ PBLs may have specific vulnerabilities versus other NHL subtypes and virus-negative tumors.

**Figure 3.**
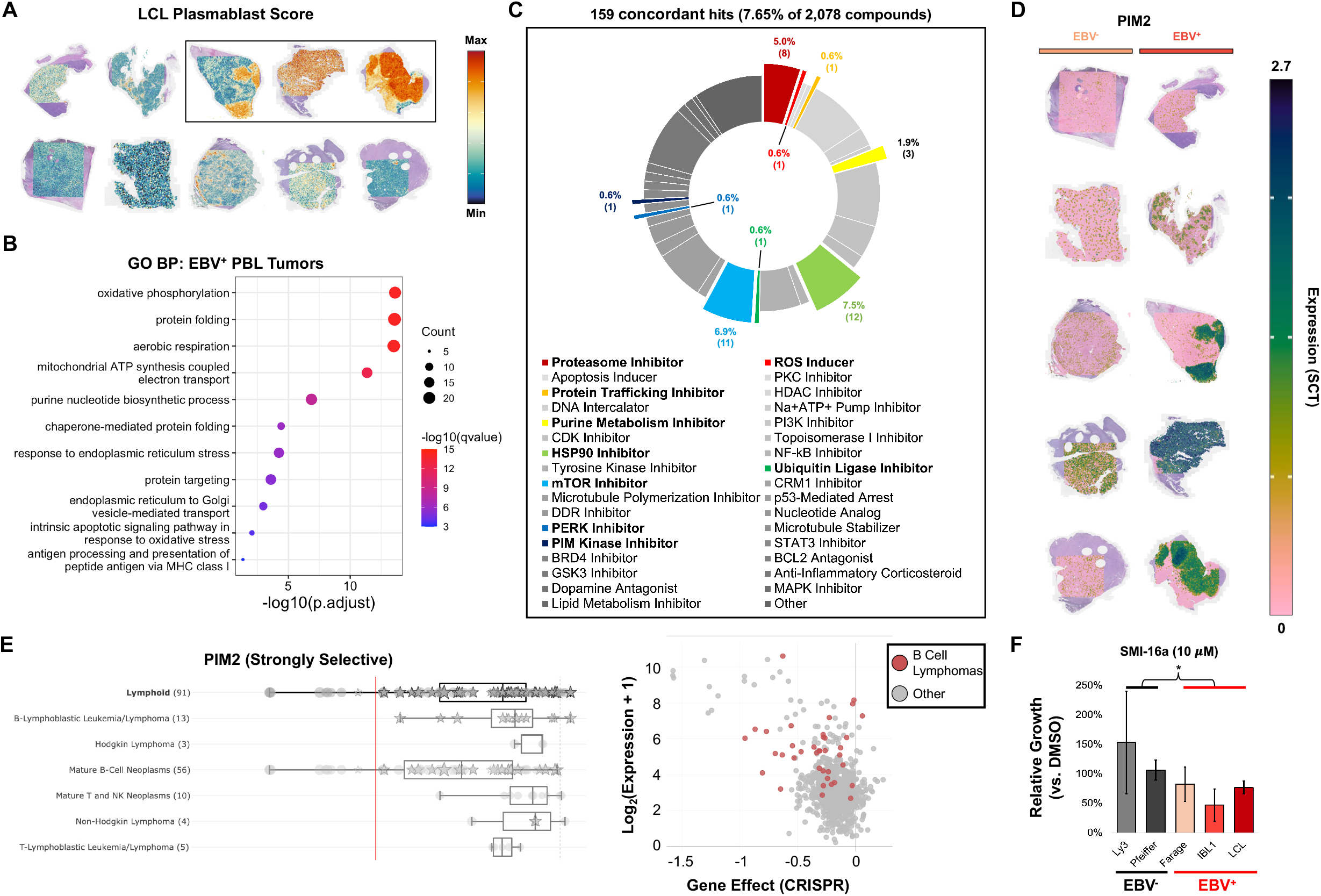
LCL growth inhibitors target pathways enriched in EBV^+^ PBLs. **(A)** HIV-NHL *in situ* plasmablast signature scoring derived from EBV-transformed LCL scRNA-seq data. EBV^+^ non-GC samples are outlined in black (from left to right: EBV^+^ immunoblastic lymphoma, EBV^+^ plasmablastic lymphoma, and EBV^+^ high grade B cell lymphoma). **(B)** Gene ontology biological process (GO BP) enrichment in the three non-GC (differentiated, plasmablast-like) EBV^+^ tumors versus other HIV-NHL tumors. **(C)** LCL small molecule inhibitor screen hits by mechanism of action (highlighted targets consistent with GO BP). The percentage and absolute number (parenthetical) of molecules with concordant growth inhibition effects are presented. **(D)** ST expression of *PIM2*, a putative target identified by LCL growth inhibition screening, in EBV^-^ and EBV^+^ HIV-NHLs. **(E)** DepMap data for PIM2 CRISPR perturbation sensitivity of 91 lymphoid lines stratified by neoplastic subtype (left panel); PIM2 CRISPR effects and expression in B cell lymphomas (red) versus other DepMap cell lines (right panel). **(F)** Sensitivity of EBV^-^ (Ly3, Pfeiffer) and EBV^+^ (Farage, IBL1, LCL) DLBCL model cell lines to a PIM2-selective inhibitor (SMI-16a, 10μ;M; n = 3 per line); statistical significance evaluated using two-tailed t-test (*p=0.039).

### Expanded EBV antigen expression at tumor-immune borders

We examined intratumoral heterogeneity across EBV^+^ HIV-NHLs in greater detail based on EBV-induced cell fates from scRNA-seq^28,31^. Three states observed in previous *in vitro* studies scored highly in EBV^+^ HIV-NHLs: B Prolif (centroblast-like), AP (pre-GC activated precursor), and Plasmablast (**Figure 4A**). The BL tumor scored uniformly for B Prolif, while the polymorphic B-LPD exhibited similar frequencies of B Prolif and AP phenotypes. A progressive shift from B Prolif to Plasmablast states was evident across immunoblastic (IB), plasmablastic (PB), and PC NHLs, indicating B cell differentiation. We examined viral expression (EBNA2, LMP1, and Zta/BZLF1) in the context of spatial tumor and immune signatures (**Figures 4B-F, S6**). As expected, BL was predominantly latency 0/I (EBER^+^; EBNA2^-^LMP1^-^BZLF1^-^). Notwithstanding, rare cells expressing EBNA2 but not LMP1 or BZLF1 – matching latency IIb – were identified. Intriguingly, EBNA2^+^ cells were near tumor-immune borders (**Figure 4B**, red ROI) and tumor-infiltrating CD14^+^ macrophages (**Figure 4B**, green ROI), though not in regions with low immune infiltration (**Figure 4B**, black ROI). Most polymorphic B-LPD tumor cells were latency 0/I; however LMP1^+^EBNA2^-^ cells (latency IIa) were present in CD4^+^ T cell-infiltrated regions (**Figure 4C**, blue and orange versus black ROI). Though anti-LMP1 signal was detected near immunoblastic HIV-NHL tumor-immune borders (**Figure 4D**, yellow and cyan ROIs) and absent from the tumor core (**Figure 4D**, black ROI), nuclear localization indicated non-specific debris staining. Only the plasmablastic NHL exhibited Zta^+^ cells, which were dispersed throughout the tumor (**Figure 4E**, magenta ROI). Higher Zta^+^ frequency coincided with the highest immune cell density (**Figure 4E**, purple ROI), and fewer Zta^+^ cells were present in dense tumor (**Figure 4E**, black ROI). One EBV^+^ HIV-NHL (HG PC lymphoma) was uniformly EBNA2^-^LMP1^-^BZLF1^-^ (**Figure 4F**). Qualitatively, these data underscore viral expression heterogeneity and provide evidence for EBV expanded latency or reactivation at tumor-immune borders.

**Figure 4.**
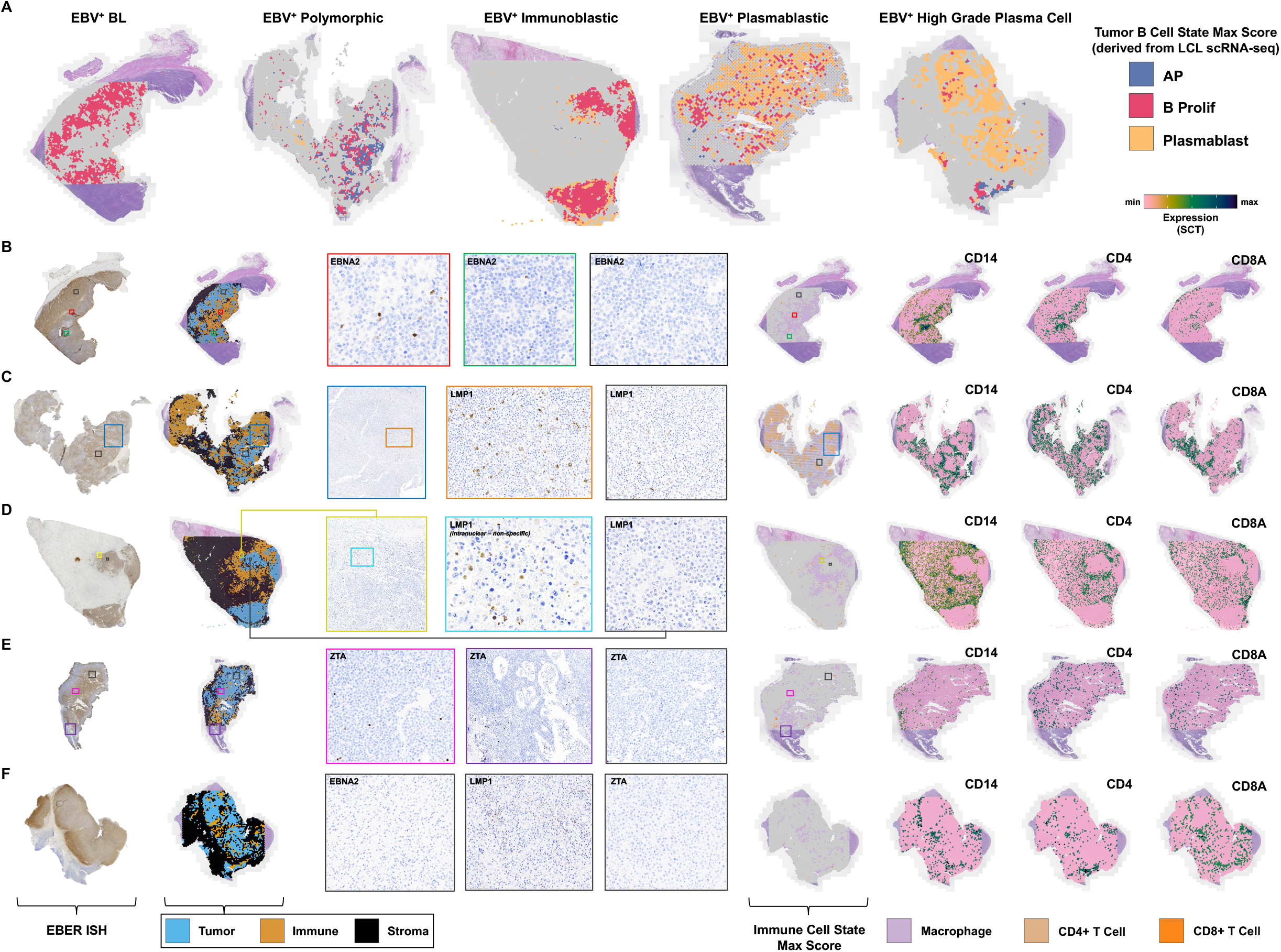
EBV^+^ cells in expanded latency and reactivation near tumor-immune borders. **(A)** ST spots exhibiting max scores for tumor B cell phenotypes (AP = activated precursor; B Prolif = actively cycling B cell; Plasmablast) in EBV^+^ HIV-NHL samples (left to right: Burkitt Lymphoma, Polymorphic B-LPD, Immunoblastic DLBCL, Plasmablastic Lymphoma, and High Grade Plasma Cell Lymphoma). **(B)** Detail of EBV EBNA2 IHC staining in BL in context of tissue segmentation and immune signatures. From left to right: EBER stain; tissue segmentation; EBNA2 staining at tumor-immune border (red box), proximal to tumor-infiltrating macrophages (green box), and in tumor core (black box); spots with maximal macrophage and T cell subset scores; and expression of macrophage (*CD14*) and T cell (*CD4, CD8A*) lineage markers. **(C)** Detail of EBV LMP1 IHC staining in polymorphic lymphoma as shown in B. Regions with high T cell density (blue and orange boxes) versus tumor cores with low T cell density (black box) are presented. **(D)** Detail of EBV LMP1 IHC staining – considered to be non-specific due to lack of membrane localization – in immunoblastic DLBCL as shown in B and C. Regions at tumor-immune border (yellow and cyan boxes) versus tumor core (black box) are presented. **(E)** Detail of EBV ZTA (BZLF1) IHC staining in plasmablastic DLBCL as shown in B-D. Regions near tumor-immune borders (magenta and purple boxes) versus tumor core (black box) are presented. **(F)** Detail of EBV EBNA2, LMP1, and ZTA IHC staining in high-grade plasma cell lymphoma as shown in B-E; negative for tested EBV proteins (black boxes).

### HIV-NHL B cell state reinforcement by microenvironment signatures

Given COO-dependent HIV-NHL stromal and immune signatures, we investigated whether HIV-NHL microenvironments recapitulated aspects of 2° lymphoid organization and microenvironmental variations correlated to B cell development. As in 2° lymphoid tissues^78,79^, *CXCL12* (SDF1) and *CXCL13* displayed inverse localization, with *CXCL12* almost exclusively in stromal cells (**Figure 5A-B**). Though we found no clear associations between EBV status and *CXCL12* or *CXCL13*, upregulated *CXCR4* expression (a CXCL12 receptor) in EBV^+^ PBL coupled with stromal *CXCL12* was noteworthy since this interaction supports PC migration, retention, and survival in bone marrow^80^ (**Figure S7**).

**Figure 5.**
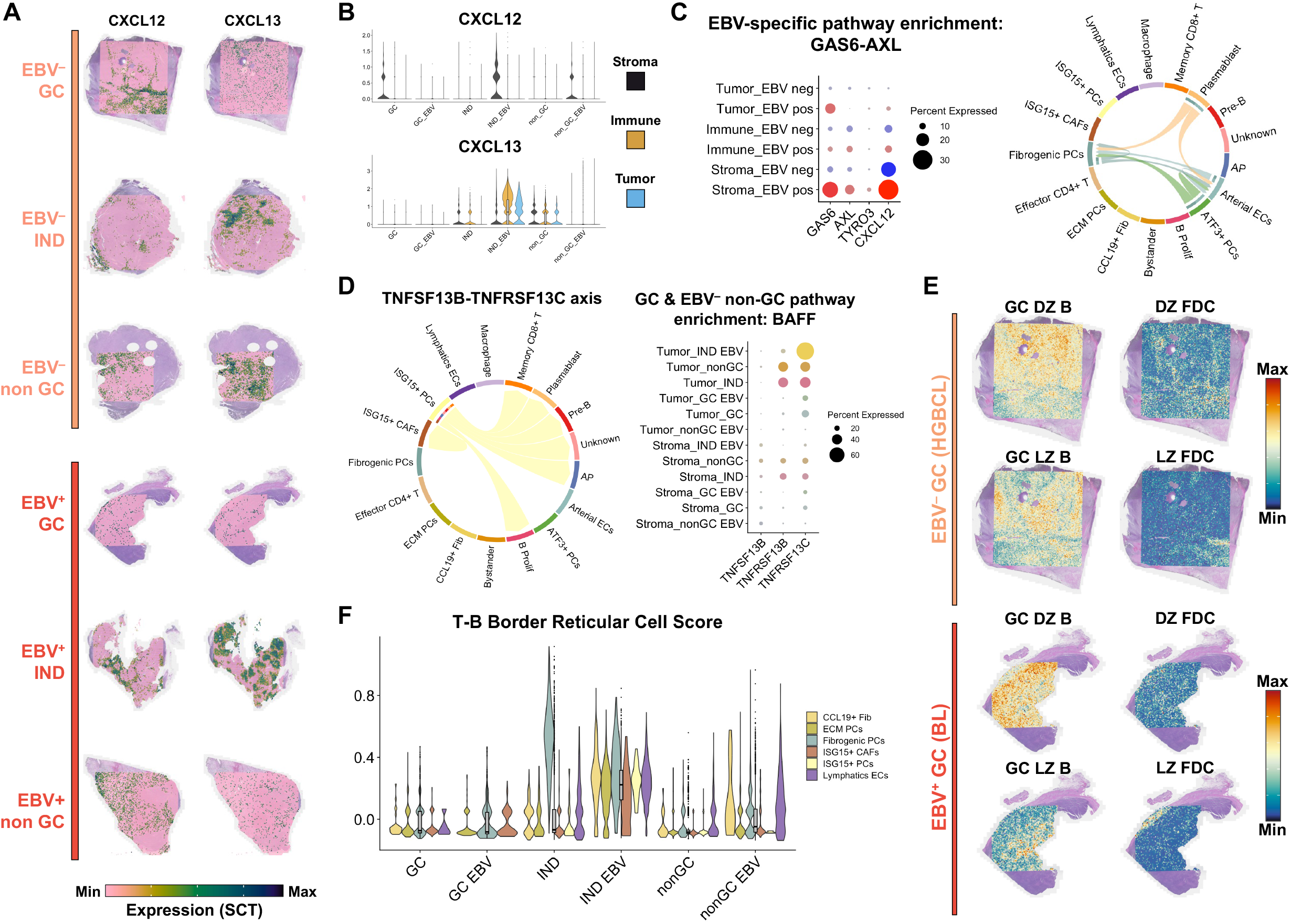
HIV-NHL TMEs reflect lymphoid compositions that reinforce tumor B cell stage. **(A)** *CXCL12* and *CXCL13* expression *in situ* within select EBV^+^ and EBV^-^ HIV-NHLs by COO. **(B)** Tissue-stratified *CXCL12* and *CXCL13* expression grouped by HIV-NHL COO and EBV. **(C)** Stratified expression of *GAS6*, TAM-family receptors (*AXL, TYRO3*), and *CXCL12* within tissues across EBV^+^ (red) and EBV^-^ (blue) HIV-NHLs (left panel); EBV^+^ HIV-NHL enrichment of *GAS6*-*AXL* cell-cell communication network with sending and receiving phenotypes denoted by chord arrows (right panel). **(D)** Enrichment of *TNFSF13B*-*TNFRSF13C* (BAFF-BAFFR) interactions in GC and EBV-negative non-GC HIV-NHLs (left panel); dotplot of BAFF pathway genes in tumor and stromal tissue stratified by COO and EBV status (right). **(E)** Expression signature scores for GC B cell states (DZ = dark zone; LZ = light zone) and follicular dendritic cell (FDC) states (DZ and LZ) in GC-derived HIV-NHLs. **(F)** Scoring of T-B border reticular cell (TBRC) expression signature within stromal phenotypes of COO- and EBV-stratified HIV-NHLs.

*CXCL12* expression is associated with GAS6-AXL interactions^81^, an axis uniquely enriched in EBV^+^ HIV-NHLs based on CellChat^64^ analysis (**Figure 5C**). *GAS6* was expressed by PBL tumors and stromal pericytes (Fibrogenic PCs & ATF3+ PCs) and predicted to interact with *AXL* (and the related receptor *TYRO3*) expressed by Fibrogenic PCs and endothelial cells. *GAS6*-*AXL* activity correlates with enhanced cell survival, angiogenesis, and metastasis in several solid tumors^82-85^ and multiple myeloma^86^.

Conversely, the BAFF-BAFFR (*TNFSF13B*-*TNFRSF13C*) axis that regulates B cell activation was enriched in GC-derived and EBV^-^ non-GC HIV-NHLs but absent in EBV^+^ PBL (**Figure 5D**). This was evident from negligible *TNFSF13B* (*BAFF*) expression in EBV^+^ non-GC stroma and lack of *TNFRSF13C* (and *TNFRSF13B* / *TACI*) expression in EBV^+^ non-GC tumors. While reduced *TNFRSF13C* expression is expected from the B cell-to-PC transition^87-89^, concomitant absence of stromal *TNFSF13B* suggests that HIV-NHL microenvironments may reflect COO-matched growth and survival conditions.

We also calculated GC B cell and FDC signatures^15^ (**File S4**) to explore 2° lymphoid aspects of tumor-microenvironment interactions. LZ and DZ B signatures were spatially distinct in EBV^+^ and EBV^-^ GC-derived HIV-NHLs (**Figure 5E**). LZ scores were higher near tumor edges and immune infiltrates. Within GC-derived HIV-NHLs, DZ B and FDC signatures were more frequent than LZ counterparts, indicating dendritic/reticular correspondence to tumor COO. The signature for extrafollicular T-B border reticular cells (TBRCs)^16^ was reduced in GC-derived HIV-NHL stroma and enriched in IND and non-GC tumors comprising differentiated B cells (**Figure 5F**). TBRC-associated expression was higher in EBV^+^ samples by COO, particularly in CCL19+ Fibroblasts, extracellular matrix pericytes (ECM PCs), and lymphatic endothelial cells (Lymphatics ECs), consistent with TBRC support of B-T interactions during post-GC differentiation prior to lymphoid egress. Thus, HIV-NHL microenvironments appear synchronized to tumor B cell phenotypes and retain stage-dependent lymphoid reticular milieus^16^.

### HIV-NHL spatial gradients reveal differential macrophage polarization dependent on tumor proximity and EBV status

We developed an approach to extract annotated spot coordinates and calculate inter-tissue distance statistics (**Figure 6A**). Spots were represented by proximity to tissue of interest, such as nearest tumor (**Figure 6B**), to identify expression gradients, spatially correlated genes (**Figure 6C**), and inter-cell proximity (**Figure 6D**). For example, we re-examined *SLC40A1* enrichment in EBV^+^ cases (**Figure 6E**). *SLC40A1* was elevated in macrophages from EBV^+^ HIV-NHLs – especially macrophages closer to tumor (**Figure 6E**) – consistent with context-dependent macrophage polarization plasticity^90^. COO- and EBV-stratified gradients of other polarization-associated genes^91^ found EBV^-^ HIV-NHL macrophages skewed toward inflammation-associated profiles (*GBP5, CYBB, TNF*) while EBV^+^ HIV-NHLs displayed immunosuppressive macrophage phenotypes (*DHCR7, A2M, SLC40A1*) (**Figure 6F**). As for *SLC40A1*, most polarization markers exhibited higher expression closer to tumor. Thus, gradient analysis between segmented tissue compartments was able to reveal distinct macrophage responses – with implications for resolving versus chronic inflammation – linked to HIV-NHL EBV status. More generally, this spatial gradient methodology provides a powerful and flexible approach to analyze distance-dependent expression between annotated phenotypes in ST datasets.

**Figure 6.**
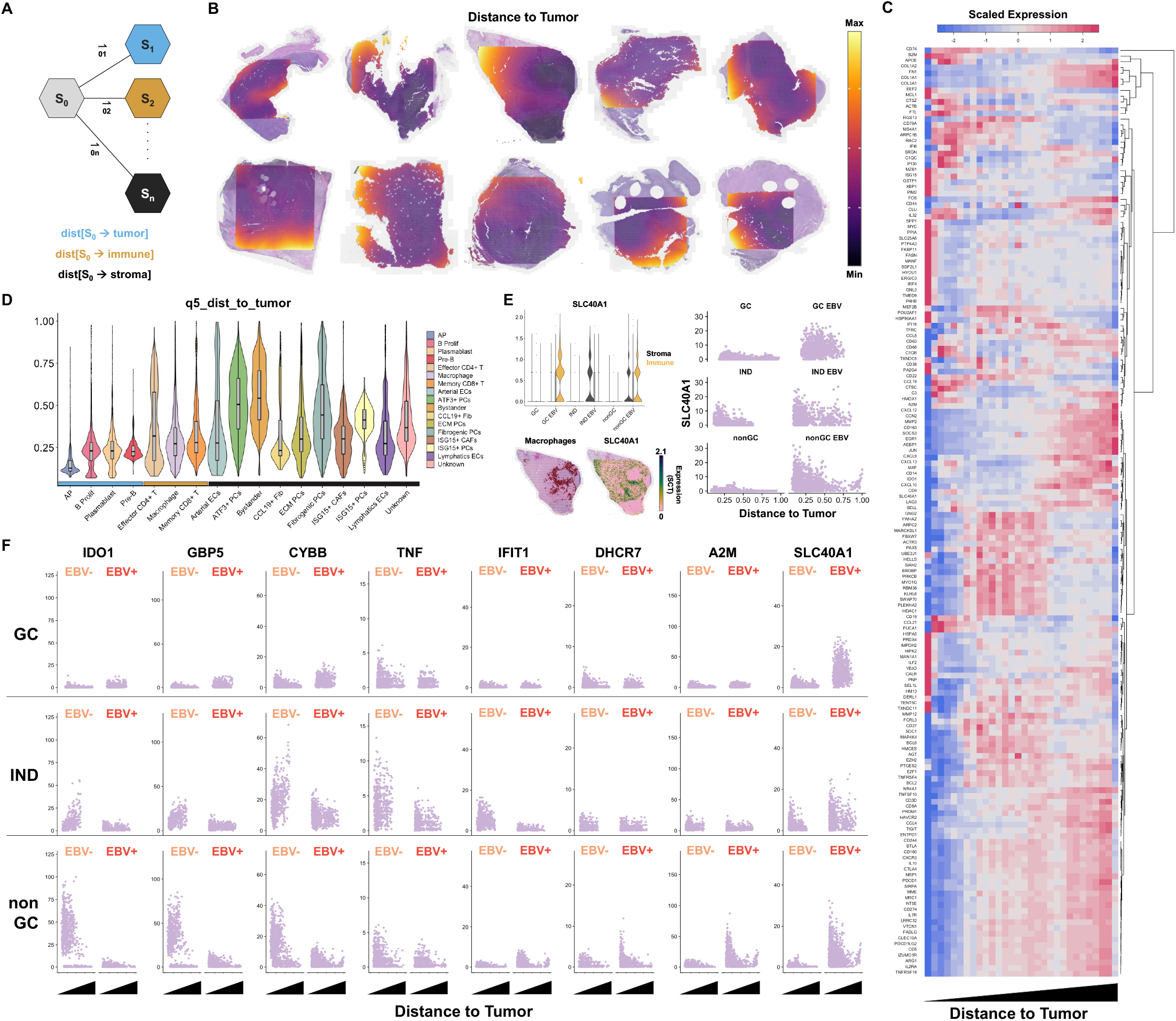
Spatial gradient analysis of HIV-NHL TME composition and expression. **(A)** Schematic of approach for calculating annotated spot-to-spot distance distributions from spatial coordinates. **(B)** ST spot representation by distance to nearest tumor region (values show the 5^th^ percentile value for each spot’s distance distribution to all annotated tumor spots: q5_dist_to_tumor). **(C)** Unsupervised clustering of spatial expression gradients by tumor proximity (scaled average expression shown in 30 bins by relative distance to tumor). **(D)** Tumor proximity distributions of ST spots by annotated cell and tissue phenotypes. **(E)** *SLC40A1* expression in immune and stromal regions (violin plot), correlation with macrophages (spatial plots), and macrophages-specific *SLC40A1* distance gradients stratified by COO and EBV status (scatter plots). **(F)** COO and EBV-stratified expression gradients of genes associated with distinct macrophage polarization. Genes are approximately ordered from left to right based on associations with M1-like (left) or M2-like (right) polarizations.

## Discussion

We present ST analysis of NHLs occurring in PLWH, including those with and without EBV. These include DLBCLs from GC and non-GC subtypes, BL, and polymorphic B-LPD. Because DLBCLs don’t always fit into a specific category in HIV and/or EBV infection contexts, cases of indeterminate COO were also included. While diverse, these cases are representative of clinical presentations; this relatively small set contains rich data that reveal important tumor-TME interactions. Specifically, our data highlight several interactions dependent on tumor B cell stage and/or EBV status that may affect oncogenicity and immunomodulation.

CD44 expression and signaling appear dependent on COO and may be influenced by EBV. CD44 is a ubiquitous glycoprotein that binds hyaluronic acid, osteopontin (OPN; *SPP1*)^92^, and collagen^93^. CD44-OPN interaction increases chemotactic responses in CD44-transfected and naturally-expressing cells^92^. Two main CD44 isoforms, (CD44s and CD44v6), are associated with cancer progression^94,95^. CD44s expression in DLBCL is significantly correlated to anatomical sites of tumor involvement and has significant prognostic value for poor outcomes in people with localized nodal DLBCL. Prognostic power increase when CD44s expression is included with the International Prognostic Index (IPI) confirms CD44 expression as an adverse predictor of occult dissemination^61^. Our data showed CD44 and *SPP1* overexpression in non-GC EBV^+^ cells and increased *SPP1* expression in surrounding stroma, which supports activation of the CD44-OPN axis in primary nodal localized tumors but simultaneously indicates opportunities for dissemination within lymph nodes. Although CD44 expression differs between nodal and extranodal DLBCL^61^, to our knowledge this is the first evidence for CD44-OPN activity correlated to COO and EBV. *CD44* expression in CD8^+^ T regions of non-GC EBV^+^ HIV-NHLs is interesting, since cytotoxic T cell CD44 interaction with tumor-derived OPN provides an immunosuppressive checkpoint blunting CTL activation and promoting immunoevasion^67^. Other studies are needed to confirm this in non-GC EBV^+^ lymphomas.

Enriched tumor-stromal GAS6-AXL interactions in EBV^+^ PBLs are intriguing. *GAS6* expression is highest in EBV^+^ IB stroma but also expressed in EBV^+^ PB and PC tumors. *GAS6* elevation is conceivable in terminally differentiated PBL tumors, as it is observed in multiple myeloma^86^. We speculate that EBV presence and acquired mutations may enhance GAS6-mediated malignancy in differentiated B cells (prolonged survival and migration^81,82,85^) by maintaining cells’ proliferative capacity. Interestingly, EBV^+^ PB and PC cases with *GAS6*-*AXL* activity display strong *CXCL12* expression associated with this axis in other tumors^81^. Intratumoral *CXCR4* expression in these cases is consistent with PC differentiation and suggests integration of GAS6-AXL and CXCL12-CXCR4 axes with respect to tissue homing, the latter being essential for PC migration to bone marrow^65,80^.

HIV-NHL tumors and TME reflect lymphoid tissue characteristics including inverse CXCL12 and CXCL13 gradients. This homology extends to COO-dependent microenvironmental matching of B cell activation signals (e.g., BAFF-BAFFR in non-PBLs) and bidirectional matching of reticular cells with tumor B cells (GC FDC biomarkers in GC-derived NHL stroma, enriched TBRC profiles in non-GC-derived NHLs). In this regard, the HIV-NHL data are partially consistent with observations from studies of 2° lymphoid organization^15,16^ and DLBCL microenvironments^13,96^; however, lymphocyte-reticulocyte interactions in NHLs are clearly perturbed versus normal lymphoid tissues with respect to inflammation, immune cell exclusion, and other pathogenic features^97^. A prior study developed a 25-gene signature for microenvironments from 4,580 DLBCL public samples and 75 original cases, which classifies four DLBCL microenvironments: “GC-like”, “mesenchymal”, “inflammatory” and “depleted”. The inflammatory category encompasses a high proportion of ABC-DLBCL cases and exhibits increased macrophage and CD8^+^ T cell frequencies^96^. This agrees with our observations that EBV^+^ tumors are mostly non-GC and exhibit M2-like immunosuppressive enrichment. Although macrophages are typically M1-like in the inflammatory category, other studies have reported M2-like macrophages in DLBCL^98^. ST and IHC correlation showed that these macrophages are not tumor-infiltrating but rather relegated to tumor edges. Thus, macrophages might physically impede anti-tumor immunity or tumor-immune communication. Accordingly, the observed distance dependence of polarization biomarker expression is particularly intriguing. The “depleted” category characterized by minimal presence of microenvironment cells and proliferating cells has been associated with BL. However, our BL sample exhibits substantial proliferating cells, macrophages, and some CD4^+^ T cells. Interestingly, increased proportions of M2-like macrophages have been previously reported in BL TMEs (independent of EBV), and a role for EBV in promoting tolerogenic TME immune conditions has been proposed^99^. Our findings that macrophage polarization biomarkers are enriched with proximity to tumor and that M1-like versus M2-like phenotype enrichment is EBV-dependent likewise suggest virus-associated pro-tumorigenic TME conditioning.

Polymorphic B-LPDs in immunodeficient people are well recognized in transplantation contexts and increasingly recognized and accepted in PLWH. They are defined as heterogenous lymphoid infiltrates with variable B, T, and histiocyte frequencies^100^. B-LPD B cells may exhibit a full differentiation spectrum, as reflected in morphological diversity. Importantly, polymorphic B-LPDs are usually EBV-associated. While rare and likely underdiagnosed, they present unique opportunities to assess EBV-infected cell behaviors and interactions *in vivo*. The initial 2003 report describing polymorphic B-LPD included ten cases from PLWH and remains the largest study to date^46^. Beyond an increased frequency of CD8^+^ versus CD4^+^ T cells, limited information about HIV-associated polymorphic B-LPD microenvironments exists. We identify a mixture of activated precursor B cells and cycling B cells but almost no plasmablasts in this tumor. Consistent with previous studies, clear T cell infiltration was also observed; however CD4^+^ T cells predominated over CD8^+^ T cells. Interestingly, CD4^+^ T cells expressed the exhaustion marker PD-1. Thus, this is the first description of an immunosuppressive microenvironment in HIV-associated polymorphic B-LPD.

This study yields genome-wide *in situ* data for tumor and TME features in HIV-NHLs resolved by tissue, COO, and EBV. Vulnerabilities predicted from clinical samples – such as PIM2 in EBV^+^ cases – were validated as targets by pharmacologic screening in cell lines. Cells with heterogeneous EBV programs were identified (as previously observed^27^) and exhibited distinct proximities to immune cells. Intratumoral EBV antigen variation likely affects tumor progression, since cells expressing different infection programs may alter immune recognition or microenvironment conditioning. Overall, enhanced immunosuppressive signatures and distinct tumor-TME activities were observed in EBV^+^ HIV-NHLs, which provide a unique resource and should inform future EBV status-dependent therapies.

## Data Availability

All sequencing data will be made publicly available via NIH GEO accession at the time of acceptance.

## Acknowledgments

We wish to thank the Duke Center for AIDS Research (CFAR) for pilot funding and the Duke Molecular Genomics Core for technical support for spatial transcriptomics experiments. We thank Dr. So Young Kim and the Duke Functional Genomics Shared Resource, which provided assistance with chemical screening.

## Author Contributions

Amy Chadburn – study design, histopathology, co-wrote & edited the manuscript

Maria Montserrat Aguilar Hernandez – contributed to data analysis, edited the manuscript Joanne Dai – performed small molecule inhibitor screen

Mitra Harrison – curated phenotype annotation, analyzed ST data

Nicolas M. Reinoso-Vizcaino – co-developed LCL scRNA-seq phenotypes Ashley P. Barry – supported cell line inhibitor screens

Emily Hocke – ST library preparation and sequencing Karen Abramson – ST library preparation and sequencing Vaibhav Jain – support for ST data analysis

Cliburn Chan – edited the manuscript

Ethel Cesarman – study design, co-wrote & edited the manuscript Micah A. Luftig – study design, co-wrote & edited the manuscript

Elliott D. SoRelle – study design, funding acquisition, analyzed ST / scRNA-seq data, co-wrote the manuscript

## Ethics Statement

Samples were obtained and utilized with Weill Cornell Medicine Institutional Review Board approval.

