## Supplementary Information and Figures for "HIV-associated non-Hodgkin lymphoma tumor-microenvironment axes differ by EBV status across cellular origins"

**Title**

**Short Title**

HIV-NHL Spatial Transcriptomics

**Authors**

Amy Chadburn^1^, Maria Montserrat Aguilar Hernandez^1^, Joanne Dai^2^^, Mitra Harrison^3^, Nicolas M. Reinoso-Vizcaino^2^, Ashley P. Barry^2^, Emily Hocke^4^, Karen Abramson^4^, Vaibhav Jain^4^, Cliburn Chan^5^, Ethel Cesarman^1^, Micah A. Luftig^2^, & Elliott D. SoRelle^2,6^

**Affiliations**

[1] Department of Pathology and Laboratory Medicine, Weill Cornell School of Medicine, New York, NY, USA, 10065

[2] Department of Molecular Genetics and Microbiology, Duke University School of Medicine, Durham, NC, USA, 27710

[3] Department of Integrative Immunobiology, Duke University School of Medicine, Durham, NC, USA, 27710

[4] Molecular Genomics Core, Duke University, Durham, NC, USA, 27710

[5] Department of Biostatistics & Bioinformatics, Duke University School of Medicine, Durham, NC, USA, 27710

[6] Department of Microbiology & Immunology, University of Michigan Medical School, Ann Arbor, MI, USA, 48109

^^^Current address: Merck Research Laboratories, Boston, MA, USA, 02115

**Corresponding Author**

Elliott D. SoRelle, PhD

**Keywords**

B cell lymphoma, non-Hodgkin lymphoma, plasmablastic lymphoma, immunosuppression, HIV, Epstein-Barr virus, cell of origin, spatial transcriptomics, polymorphic lymphoproliferative disorder

**Bioinformatics Methods Detail**

Read assembly & alignment: Sequence data were assembled into sample-demultiplexed reads (.fastq) using *mkfastq*, a wrapper of *bcl2fastq* (v2.20; Illumina, San Diego, CA) using SpaceRanger (v2.0.1; 10x Genomics, Pleasanton, CA). Reads were aligned to a pre-built SpaceRanger reference genome (refdata-gex-GRCh38-2020-A) using the *count* function for the v2 probe set (--*probe-set* argument; Visium_Human_Transcriptome_ Probe_Set_v2.0_GRCh38-2020-A.csv). Slide information was provided for SpaceRanger *count* as described in standard pipelines. High resolution CytAssist (--*cytaimage* argument) and AxioScan images (--*image* argument) with array fiducials were specified to align barcoded reads. Images and QC-filtered UMI outputs (barcodes.tsv.gz, features.tsv.gz, matrix.mtx.gz) were used for downstream analyses.

Data integration and QC: UMIs and aligned images were prepared as Seurat objects in RStudio using *Read10x_Image* and *Load10x_Spatial* functions in Seurat v5^29,30^. Mitochondrial read fractions were calculated and cell cycle scores were assigned based on the “s.genes” and “g2m.genes”. In addition to default SpaceRanger QC filters, spots with <200 unique or <500 total transcripts were excluded. Sample reads were log-normalized with the *NormalizeData* function, and the top 2000 variable features were identified using *FindVariableFeatures*. Samples were then integrated via transfer anchors into a single object with *SelectIntegrationFeatures*, *FindIntegrationAnchors*, and *IntegrateData*. Integrated data were scaled (*ScaleData*), transformed (*SCTransform* with “vst” method), dimensionally reduced (*RunPCA*, *RunUMAP*), and clustered using unsupervised methods (*FindNeighbors*, *FindClusters*). Metadata labels were added to denote EBV status and COO of each HIV-NHL for DE comparisons.

Celltype annotation: Spot celltype composition was scored using scType^31^. The default scType reference was appended with expression signatures from immune^32^, cancer-associated stroma^33^, and EBV-immortalized B cell phenotypes^34^ from scRNA-seq datasets (**File S1**). Max-scored annotations were mapped to seventeen fine-grained clusters identified from unsupervised methods (sixteen known, one unknown). Fine-grained clusters were grouped to produce three coarse-grained tissue annotations: tumor (B cells in histopathologic tumors), immune (primarily T cells, NK cells, and monocytes/macrophages), and stroma (primarily fibroblasts, pericytes, and endothelial cells). Tissue spot annotations were utilized in DE comparisons. Separately, we evaluated transfer anchor integration^30^ to map phenotypes from LCL and PBMC scRNA-seq. This approach yielded similar results for tumor and immune spots but was not appropriate for stromal analysis.

Differential expression, ontology, predicted interactions and activity: DE analyses of transformed reads (SCT assay) were performed to compare samples by tissue annotation, EBV status, and/or COO using *FindMarkers* and *FindAllMarkers*. Gene set signature scores were calculated with the *AddModuleScore* function. GO enrichment was evaluated with clusterProfiler *enrichGO* for “Biological Process” (BP) terms^35^. Cellchat interaction analyses were performed for aggregated and EBV-stratified samples using scType annotations as class identities. CollecTRI and Progeny tools from decoupleR^36^ were used to calculate transcription factor and cancer-related signaling activity.

Spatial gradient analysis: Spot coordinates (row, column) were used to calculate spot-to-spot distance distribution statistics (min, mean, quantile) to quantify proximities of annotated tissue types. Relative spot distances to select tissue types were assigned as metadata, enabling expression gradient visualization informed by histology and spot annotation. Code to generate distance information is provided supplementarily (**File S2**).

**Supplementary Tables and Figures**

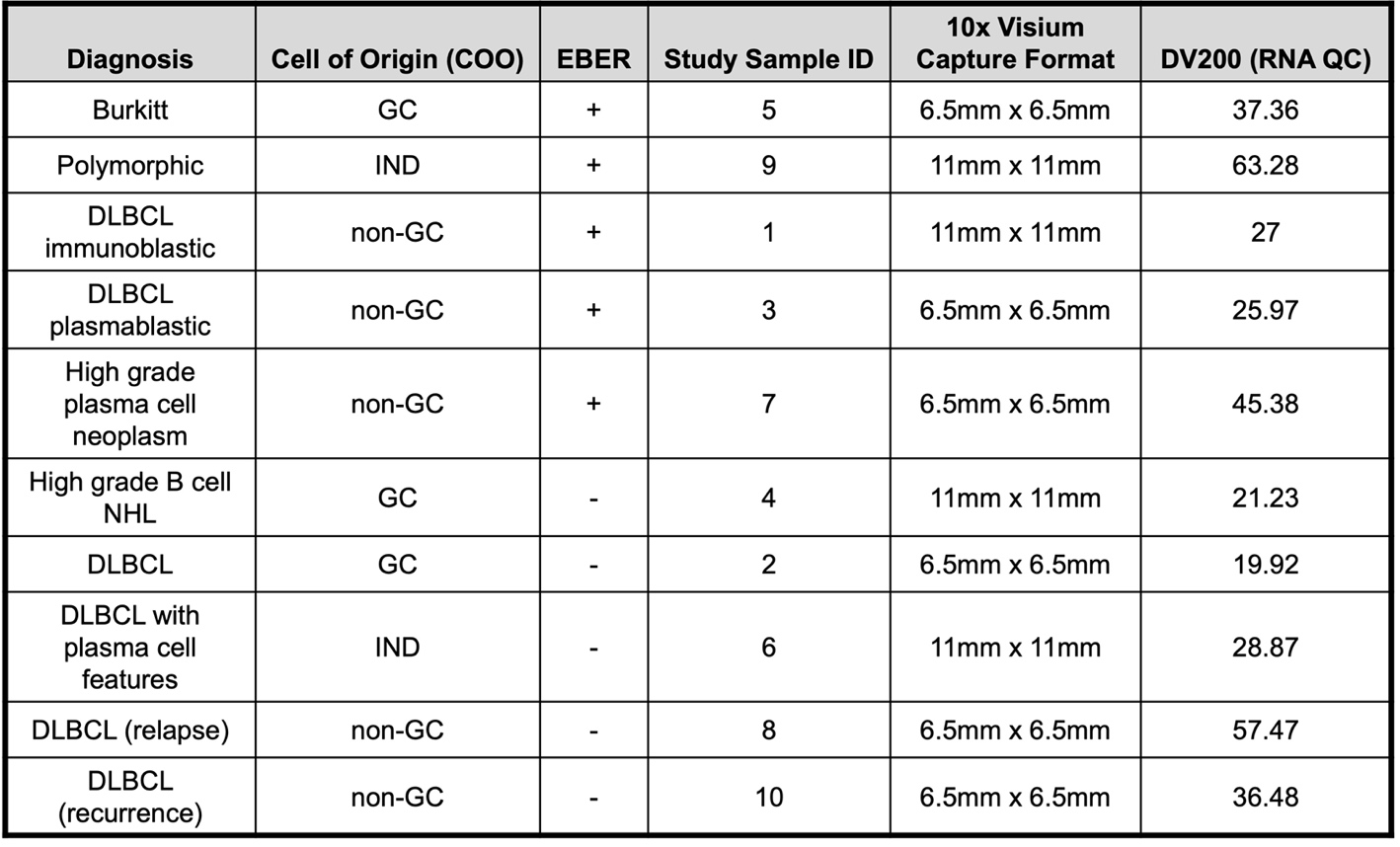

**Table S1. HIV-NHL sample overview**

Diagnostic classifications, ST capture array formats, and RNA QC values for HIV-NHL samples in this study.

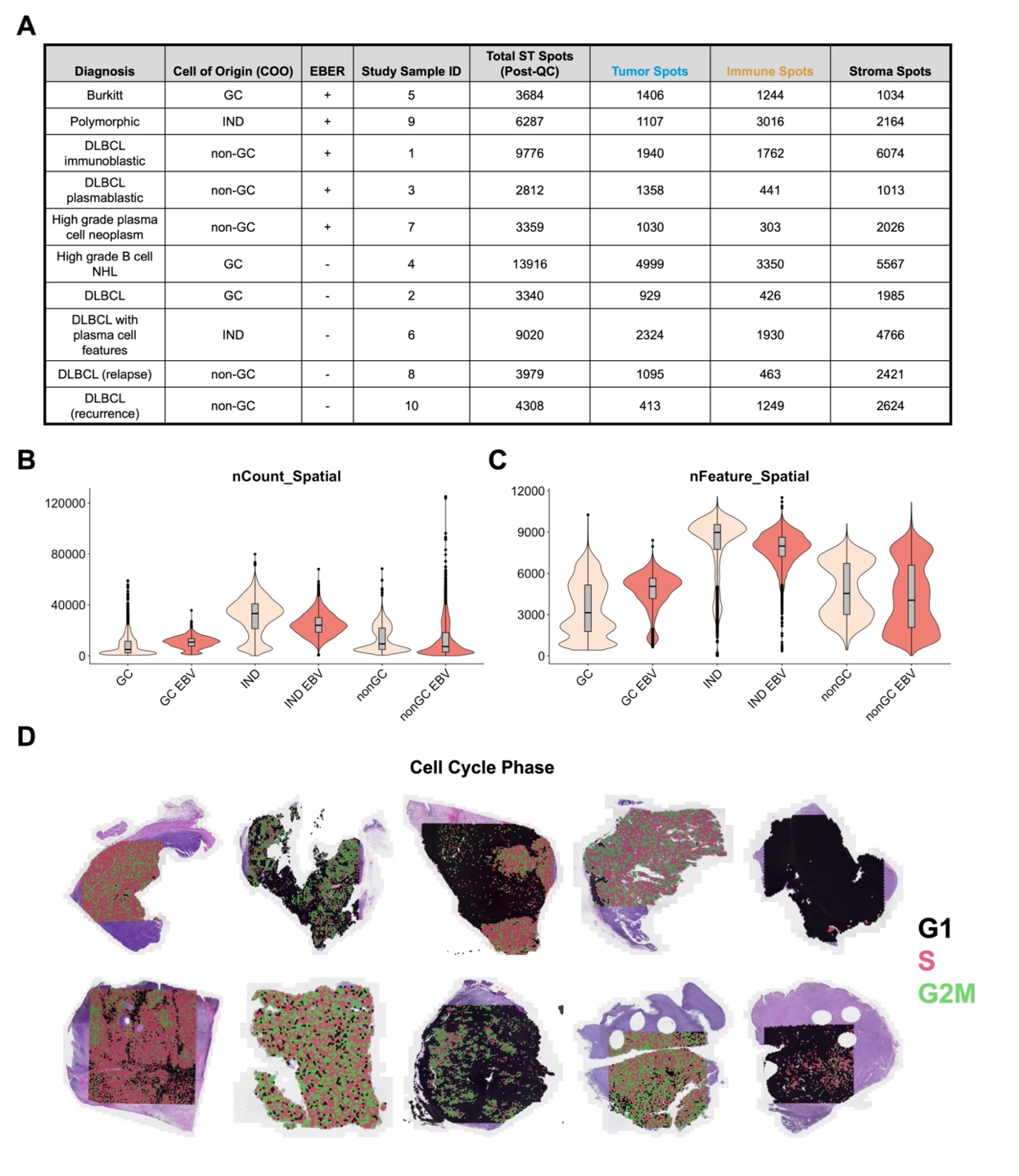

**Figure S1. QC summary of HIV-NHL ST data**

**(A)** # of ST spots by sample. Post-QC spot numbers are listed for whole samples and annotated tissue classes.

**(B)** Violin plot of total captured ST transcripts (nCount) per sample grouped by COO and EBV status.

**(C)** Violin plot of unique genes (nFeature) per sample grouped by COO and EBV status.

**(D)** Cell cycle phase annotation *in situ*.

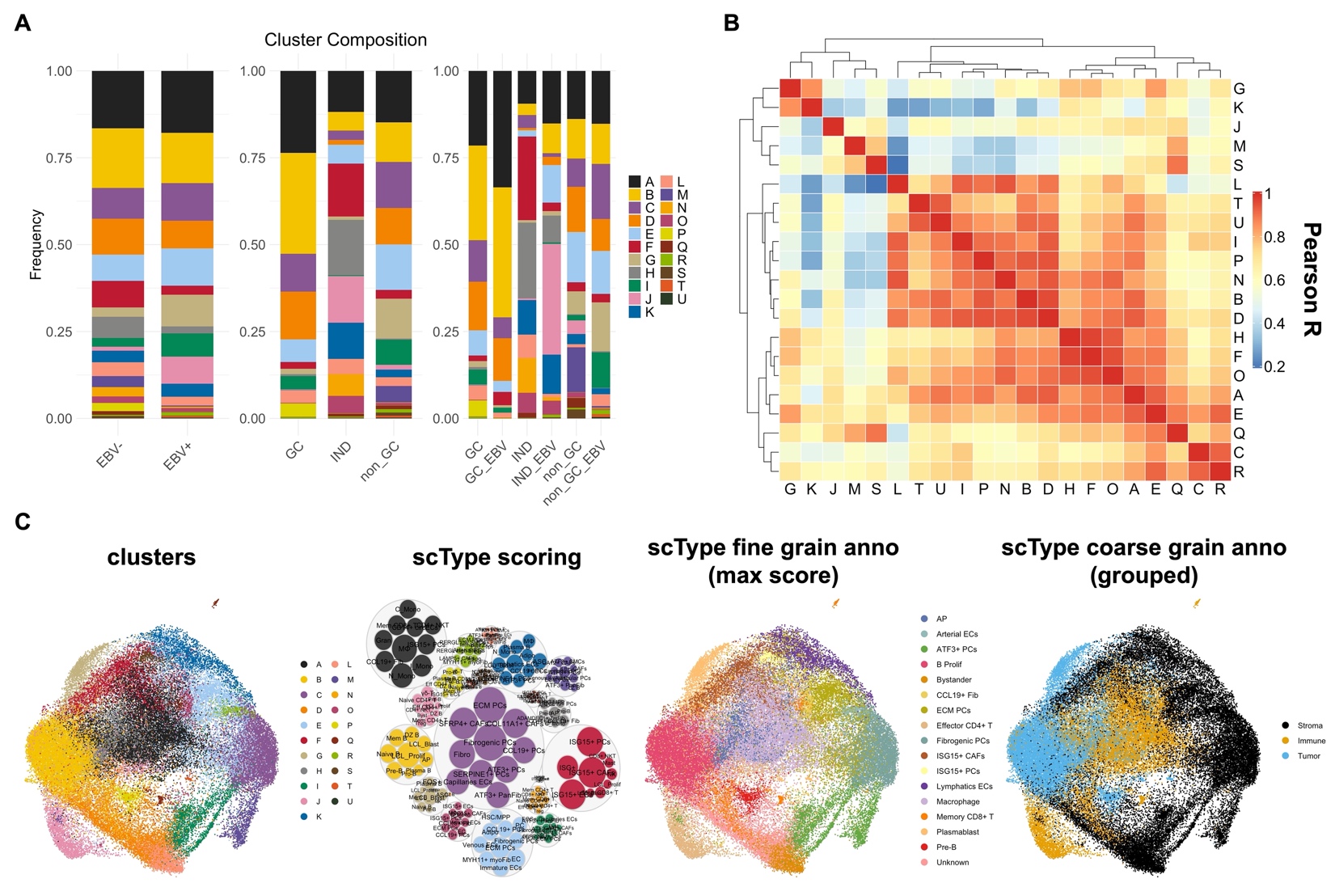

**Figure S2. HIV-NHL capture spot annotations**

**(A)** Normalized bar graphs showing unsupervised cluster compositions by EBV status (left), COO (middle), and EBV plus COO status (right).

**(B)** Pairwise genome-wide correlations (Pearson R) between unsupervised clusters.

**(C)** UMAP of ST spot clusters (far left), bubble plot of scType phenotype scoring by cluster (middle left), fine-grained scType annotation (middle right; based on max scType score per spot), and mapping to coarse-grained tissue phenotypes (far right).

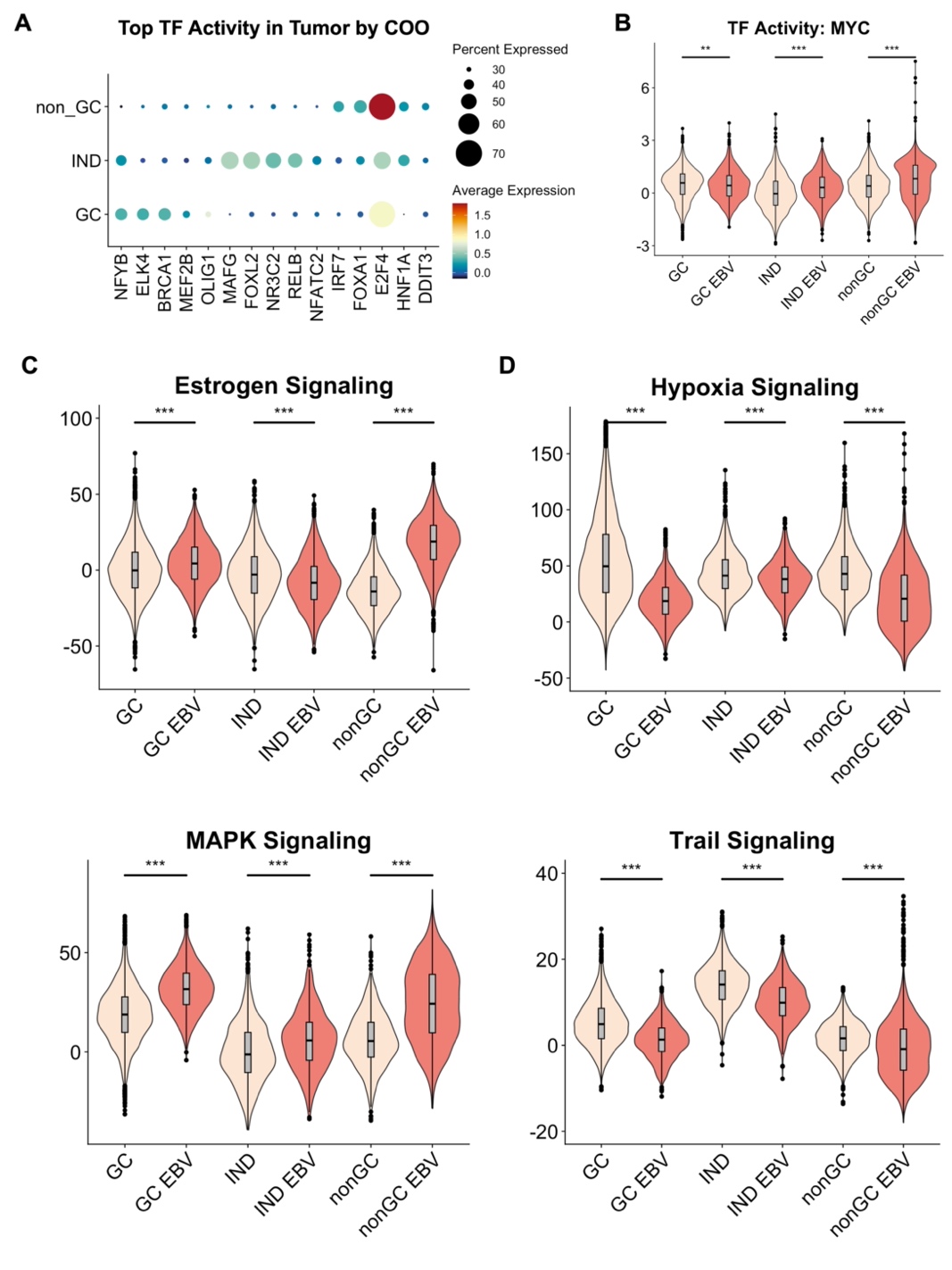

**Figure S3. Transcription factor and signaling activity inference in HIV-NHLs**

**(A)** Transcription factor activity predictions in tumor tissue based on SCT expression stratified by COO.

**(B)** Predicted MYC transcriptional activity in HIV-NHLs grouped by COO and EBV status. Asterisks denote statistically significant EBV-stratified differences in activity within COO class by non-parametric distribution test (Kolmogorov-Smirnov, **p<0.001, ***p<1e-5).

**(C)** Predicted estrogen and MAPK signaling activity in HIV-NHL tumor tissue grouped by COO and EBV status. Asterisks denote statistically significant EBV-stratified differences in activity within COO class by non-parametric distribution test (Kolmogorov-Smirnov, ***p<1e-5).

**(D)** Predicted hypoxia and TRAIL signaling activity in HIV-NHL tumor tissue grouped by COO and EBV status. Asterisks denote statistically significant EBV-stratified differences in activity within COO class by non-parametric distribution test (Kolmogorov-Smirnov, ***p<1e-5).

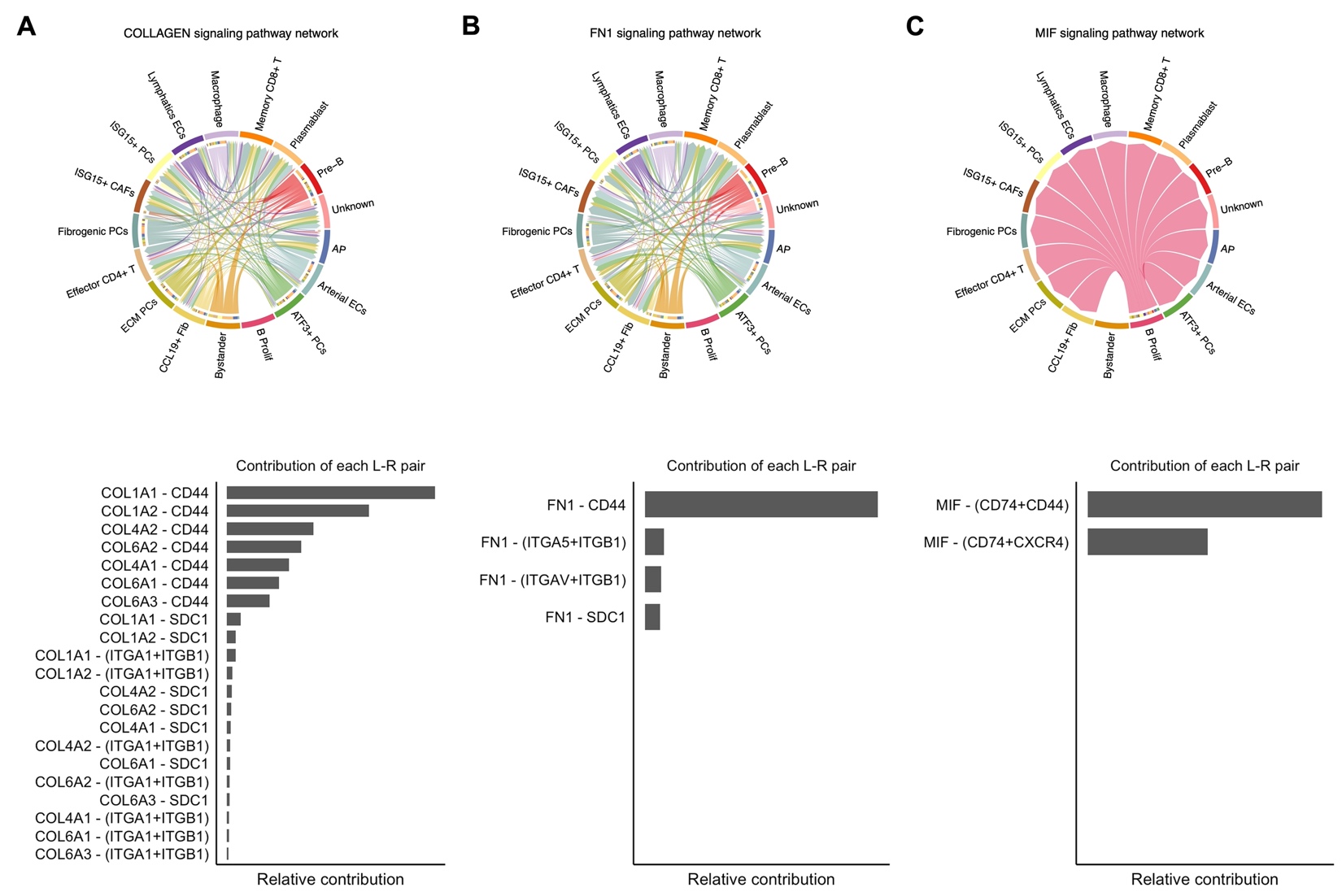

**Figure S4. Cell-cell interactions convergent on CD44 in HIV-NHLs**

**(A)** Cellchat collagen signaling network predictions across integrated HIV-NHLs. Chord colors denote phenotypes expressing ligands (senders) and arrows denote phenotypes expressing receptors (receivers). Bar plot denotes the relative contributions of specific interactions.

**(B)** Cellchat FN1 signaling network predictions across integrated HIV-NHLs, as in A.

**(C)** Cellchat MIF signaling network predictions across integrated HIV-NHLs, as in A-B.

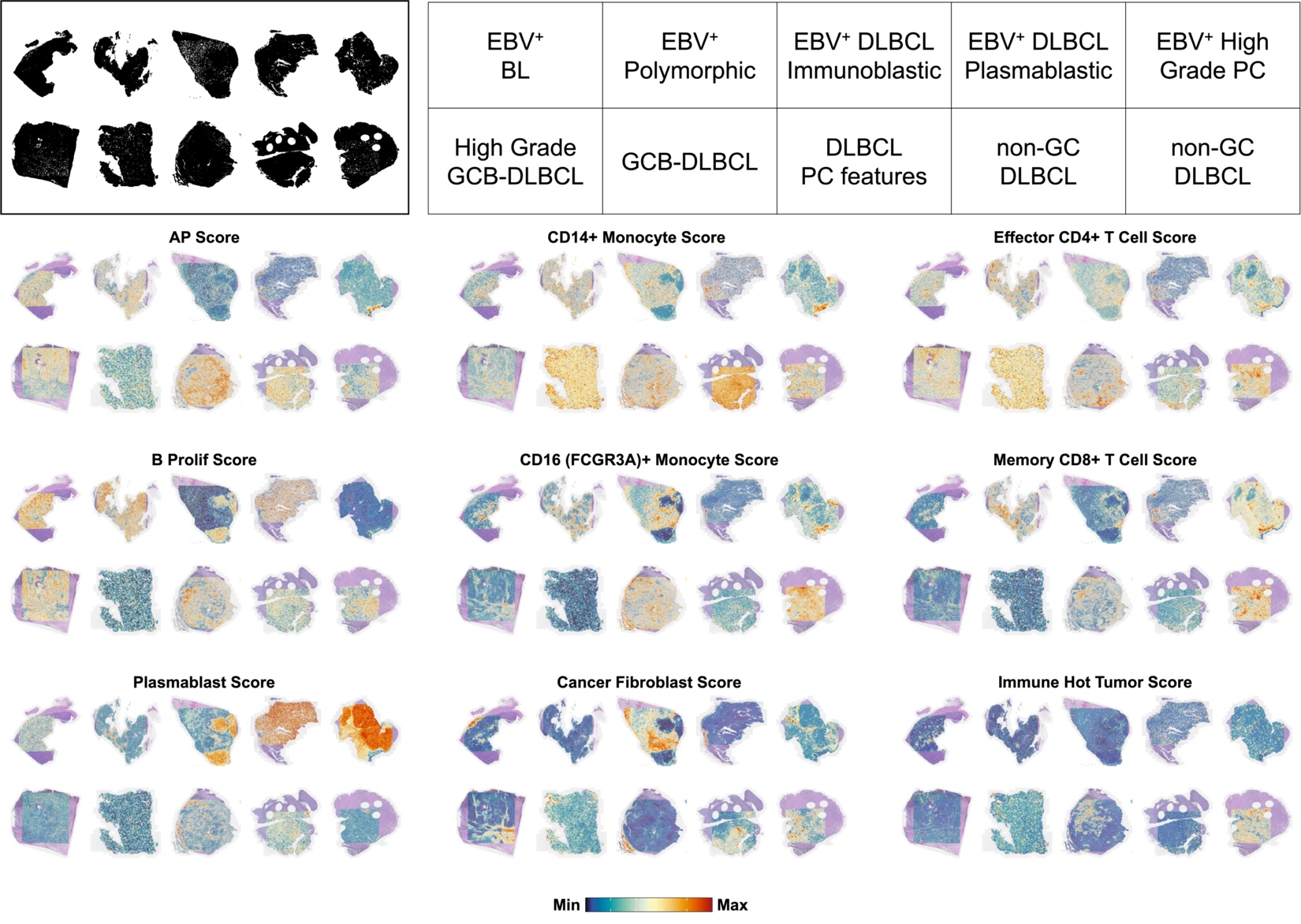

**Figure S5. Spatial scoring of LCL and immune cell scRNA-seq reference signatures**

LCL substate score spatial plots from LCL scRNA-seq reference (AP, B Prolif, Plasmablast); immune cell score spatial plots from PBMC scRNA-seq references (CD14+ Monocyte, CD16+ Monocyte, Effector CD4+ T Cell, Memory CD8+ T Cell); cancer signature modules (Cancer Fibroblast, Immune Hot Tumor).

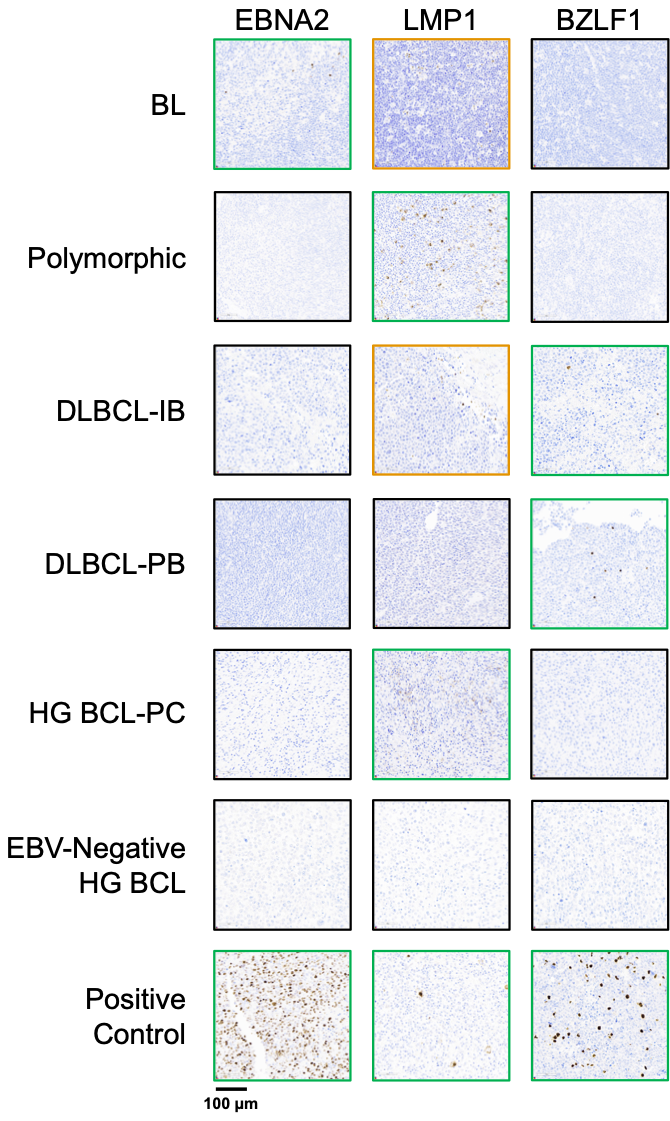

**Figure S6. Overview of viral protein IHC in EBV^+^ HIV-NHLs**

Representative images for EBNA2, LMP1, BZLF1 immunohistochemical staining across EBV^+^ HIV-NHLs (green outline = positive, gold outline = non-specific staining, black outline = negative).

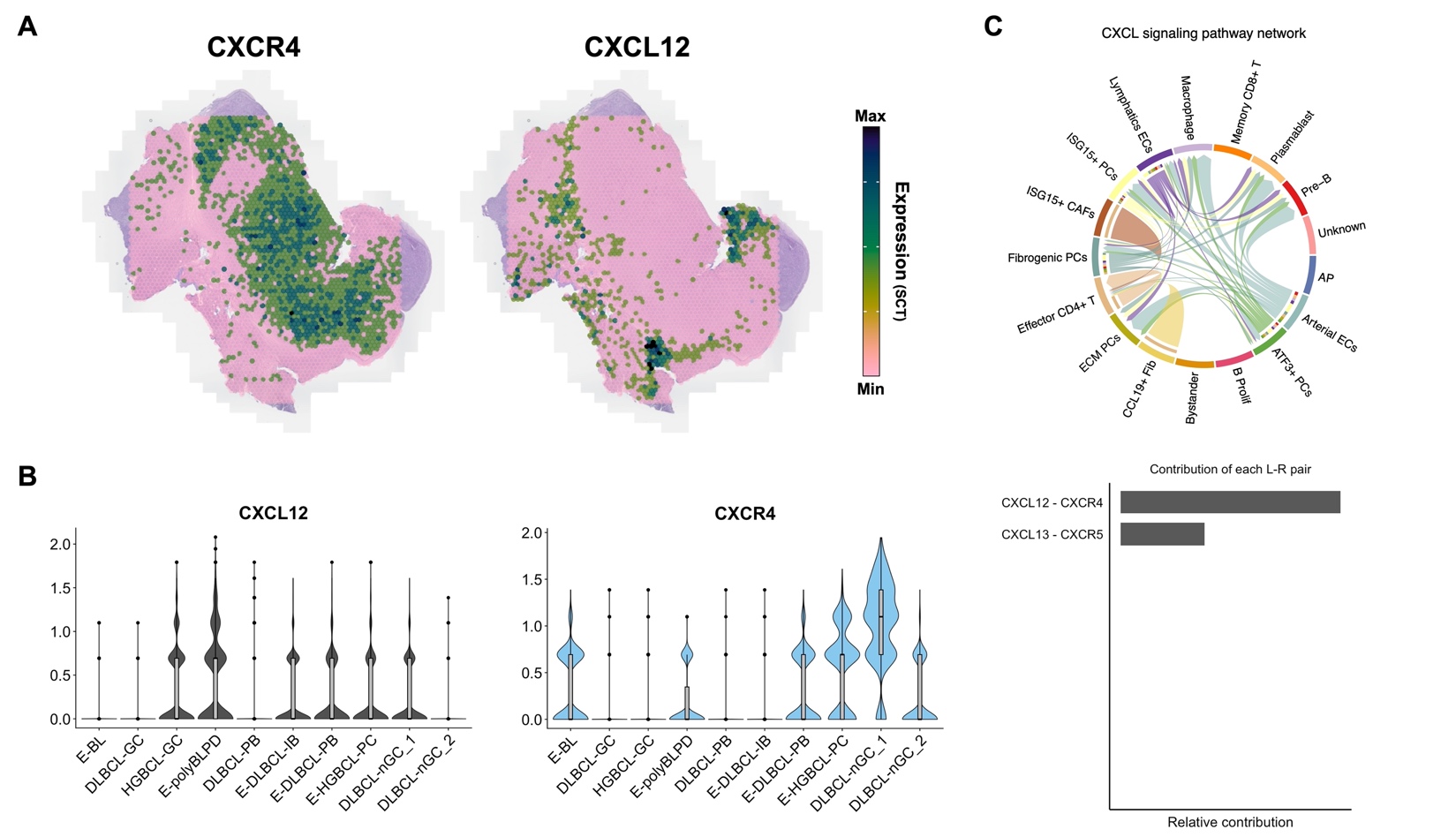

**Figure S7.** **Spatial expression of 2° lymphoid tissue chemokines and receptors in HIV-NHLs**

**(A)** *CXCR4* and *CXCL12* spatial expression in EBV^+^ HG PC NHL.

**(B)** Violin plots of *CXCL12* in stroma spots and *CXCR4* expression in tumor spots for each HIV-NHL sample.

**(C)** Cellchat analysis of CXCL family interactions (*CXCL12-CXCR4* and *CXCL13-CXCR5*).
